# Unveiling Oligodendrocyte-Lineage Differentiation in the Glioblastoma Infiltrative Zone Through Spatial Transcriptomics

**DOI:** 10.1101/2025.10.20.683359

**Authors:** Chiara Maria Argento, Enrico Sebastiani, Emanuele Filiberto Rosatti, Denise Sighel, Sara Longhi, Alessia Soldano, Toma Tebaldi, Alessandro Quattrone, Francesco Corsini, Luciano Annicchiarico, Mattia Barbareschi, Lorenza Pecciarini, Pietro Luigi Poliani

## Abstract

IDH-wildtype glioblastoma (GBM) is the most aggressive primary brain tumor, characterized by limited therapeutic options and inevitable recurrence. This recurrence is driven by the highly infiltrative behavior of GBM cells, which prevents complete resection.

Here, we performed spatially resolved transcriptomic profiling of four surgically defined GBM regions to characterize the infiltrative tumor compartment. Across multiple datasets and profiling approaches, the infiltrative zone showed reduced cellular heterogeneity compared with the tumor core. By comparison with an integrated reference of single-cell RNA sequencing data, we revealed that this reduction reflects a preferential differentiation of infiltrative GBM cells toward an oligodendrocyte-like lineage, accompanied by attenuated inflammatory signaling. In contrast, necrotic and proliferative regions display pronounced astrocyte-like and mesenchymal-like programs. Combined immunohistochemistry and fluorescence in situ hybridization confirmed the enrichment of oligodendrocyte-like tumor cells in the infiltrative area. Together, these findings identify the oligodendrocyte-like state as a defining feature and potential therapeutic vulnerability of infiltrative GBM driving brain invasion and relapse.

## Introduction

Glioblastoma IDH wild-type (GBM) is the most aggressive primary brain tumor, with only about 5.5% of Glioblastoma IDH wild-type (GBM) is the most aggressive primary brain tumor, with only about 5.5% of patients surviving for five years from the time of diagnosis^1^. Unfortunately, even after multimodal treatment, GBM inevitably recurs, with 90% of patients experiencing recurrence within two years of initial diagnosis^2^. Since recurrence is primarily due to the aggressive and heterogeneous nature of the tumor and its ability to infiltrate the surrounding brain tissue, studying GBM heterogeneity and, in particular, the infiltrative GBM cell population driving brain invasion and relapse could unravel potential therapeutic vulnerabilities and lead to the development of strategies that could effectively target the residual disease.

In recent years, several studies using single-cell RNA sequencing (scRNA-seq) technologies greatly advanced our comprehension of GBM heterogeneity, revealing that GBM cells belong to four cell identity states, which coexist in all patients^3–7^. Three of these four cell states resemble the physiological lineages of vertebrate neurodevelopment (neuronal, astrocytic, and oligodendrocytic), while the remaining state is an inflammation-related, mesenchymal-like cell state. Interestingly, several experiments have shown that GBM cells can transit between these four GBM cell states, suggesting that these states can be plastic and interchangeable^3,8–10^.

Despite their contributions, scRNA-seq studies fall short of providing the essential spatial context needed to unravel GBM heterogeneity. This spatial context is crucial for a deep characterization of the cell populations present in the different tumor areas and a nuanced understanding of the interactions between them and with the brain microenvironment. After the introduction of spatial transcriptomics (stRNA-seq) methods^11,12^, some researchers have employed this technology to start unravelling the spatial organization of GBM, identifying hypoxia as a master regulator of GBM architecture^13–18^. Nevertheless, a deep characterization and a comparative analysis of the tumor infiltrative zone in its spatial context is still lacking.

The infiltrating GBM area can be identified by T2 hyperintensity regions, and by Fluid Attenuated Inversion Recovery (FLAIR) regions in magnetic resonance imaging. However, tumor brain infiltration often extends beyond the FLAIR-hyperintense contrast-enhancing area and GBM cells can also be present in areas that may initially appear unaffected^19^. Recently, increasing attention has been directed toward this FLAIR-hyperintense, non-contrast-enhancing region, which has been recognized as clinically and oncologically significant^20,21^, given the growing evidence that most recurrences and progression occur within this area^22^. Indeed, the surgical approach for high-grade gliomas is shifting from the traditional goal of gross total resection, defined as the removal of the contrast-enhancing tumor, to a more comprehensive assessment of the extent of resection that includes the residual non-contrast-enhancing tumor^20,21^. Likewise, radiotherapy planning is being refined to incorporate these insights^22^, with increasing evidence indicating that integrating these novel surgical and radiological approaches is associated with improved overall survival^20,22^, underscoring the importance of accurately characterizing the infiltrative tumor regions where GBM cells persist after surgery.

In this study, we sampled four surgically defined and spatially distinct GBM areas and performed spatially resolved transcriptomic profiling to study tumor regional heterogeneity and characterize the cancer cells within the infiltrative zone. By combining our results with the recent findings on GBM cell states from scRNA-seq, we show that the necrotic core and the actively proliferating region of GBM exhibit high heterogeneity and are predominantly populated by mesenchymal-like and astrocyte-like cells, while in the infiltrative zone heterogeneity is reduced due to a cellular shift toward the oligodendrocyte progenitor-like phenotype. By uncovering the oligodendrocyte-like state of GBM cells as a central driver of brain invasion and disease relapse, our findings open new opportunities for therapeutic strategies aimed at disrupting lineage-specific programs to prevent recurrence and extend patient survival.

## Results

### Spatially resolved analysis of different GBM areas reveals low inter-patient heterogeneity in the infiltrative zone

To elucidate the heterogeneity of GBM cell subpopulations across distinct tumor regions and investigate the phenotypes of infiltrating glioma cells, we collected tumor tissue biopsies during surgical procedures using navigation sampling and Gliolan fluorescence. GBM samples were systematically obtained from four locations in each patient (**Figure 1A**), enabling the compilation of a high-resolution spatial transcriptomics dataset comprising 12 samples from 3 patients. Detailed patient characteristics and molecular features of the respective samples are provided in **Table S1**. For each patient, we selected four regions to ensure comprehensive coverage: a region within the tumor core that exhibits distinct histological features, primarily areas of pseudopalisading necrosis, referred to as the **core** of the tumor; an area within the tumor marked by intense cell proliferation and strong Gliolan fluorescence, referred to as the **inner edge** of the tumor; a transition area between the tumor and the surrounding healthy tissue, characterized by weak Gliolan fluorescence and FLAIR hyperintensity, indicative of infiltrative activity, hereby termed the **outer edge** of the tumor; and a **distal area** from the tumor, with supposedly healthy tissue (**Figure 1A**).

**Fig. 1:**
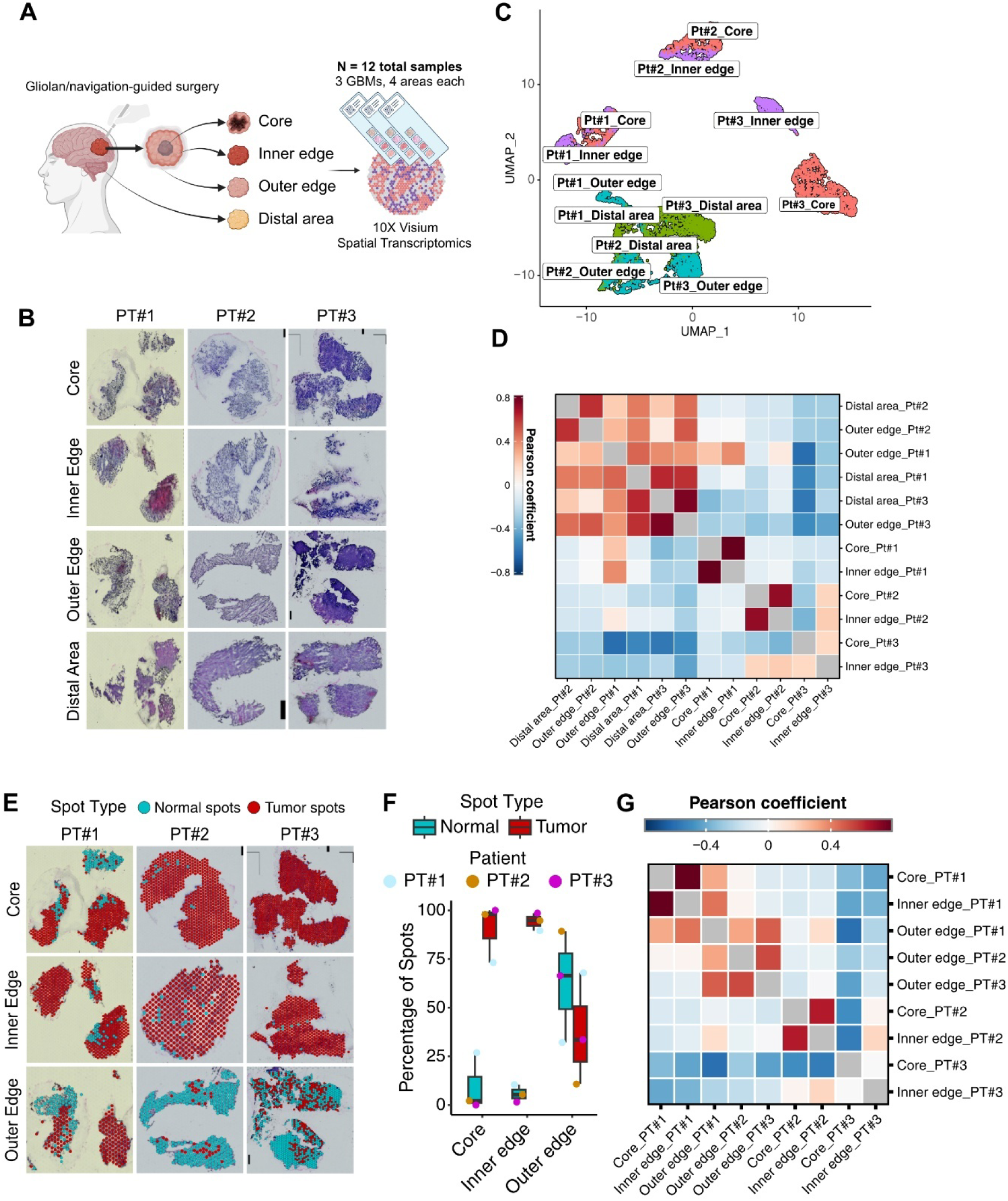
Spatial transcriptomics analysis of four different GBM areas reveals a differential profile of inter- and intra-patient heterogeneity. **(A)** Outline of the spatial transcriptomics dataset showing the sampled areas for each patient in the project. We selected four areas for sampling. The “core” and the “inner edge” are located inside the tumor. The former one is at the center and is predominantly necrotic; the latter one is the main proliferative region of the tumor. The third is the infiltrative tumor margin and the last is a distal area from the tumor, containing only healthy cells. A total of N=12 samples, comprising 4 areas for 3 different patients, were then processed with the stRNA-seq Visium protocol. **(B)** H&E staining of the four sampled areas for each patient (PT) located within the capture area of the Visium slide. **(C)** UMAP plot computed on the integrated stRNA-seq dataset after NMF reduction. Colors indicate the four different sampled areas. **(D)** Heatmap of Pearson correlation coefficients of all spots in the different areas of the three patients. Rows and columns are hierarchically clustered to highlight similarity. **(E)** Surface plots showing tumor and non-tumor spot assignment based on the CNV analysis. **(F)** Boxplots showing tumor or normal identity assignment for all spots based on the CNV analysis. **(G)** Heatmap of Pearson correlation coefficients of spots assigned as tumor in the different areas of the three patients. Rows and columns are hierarchically clustered to highlight similarity.

The resected tumor samples were embedded, cryosectioned, and stained with hematoxylin and eosin (H&E) (**Figure 1B**). The samples were then processed according to the stRNA-seq Visium protocol (**Figure 1A**, **Figure 1B, Table S2**). In the 10X Visium spatial transcriptomic platform the diameter of the capture spot is 55 mm, and each spot contains a mixture of 1–35 cells, with a median of 8 cells in GBM samples based on image analysis ^17^. Each sample had a comparable number of spots for each area (**Figure S1A**). Initial data quality was assessed by examining gene expression counts, mitochondrial gene ratios, and complexity across the spots (**Figure S1B-E**). After filtering, we retained 7433 spatial spots for downstream analysis. To visualize the distribution of sampled areas in a Uniform Manifold Approximation and Projection (UMAP), we employed a non-negative matrix factorization (NMF) reduction method (**Figure 1C**). Upon integrating all sampled areas, the outer edge of the tumor and the distal area consistently clustered together across patients. In contrast, the core and inner edge of the tumor exhibited distinct clustering patterns unique to each patient (**Figure 1C**). To further evaluate the degree of similarity among the areas, we performed linear correlation of the transcriptomes of all sampled regions across all patients. In concordance with the UMAP, the tumor core and the tumor inner edge showed no correlation between different patients (median Pearson correlation coefficient of −0.09 ± 0.34) (**Figure 1D**), although these areas exhibited higher similarities within the same patient. Conversely, there was a moderate degree of similarity between the outer edge of the tumor and the distal area in all patients (median Pearson coefficient of 0.43 ± 0.18) (**Figure 1D**).

These findings underscore that the strong inter-patient heterogeneity observed in GBM is primarily driven by the tumor bulk regions, which exhibit substantial molecular and cellular diversity. In contrast, the infiltrative margins and adjacent healthy tissues display markedly lower levels of heterogeneity, suggesting a more conserved or uniform biological profile across patients.

### Reduction of cellular complexity in the infiltrative zone suggests a common mechanism of infiltration in GBM

To better characterize the surgically defined regions of GBM, and exclude that the low inter-patient heterogeneicity of single-cell transcriptome profiles observed in the outer edge region could be attributed to a greater prevalence of non-tumor cells, we assessed the content of tumor cells in our four areas via immunohistochemistry (IHC) analysis of tissue sections near-adjacent to those used for the spatial transcriptomics protocol. We performed IHC for two common GBM markers - MKI67 and SOX2. As expected, the distal area-derived sections showed significantly lower expression for both MKI67 and SOX2 compared to the tumor core, inner edge, and outer edge (**Figure S1F**), confirming the lack of tumor cells in this area. Conversely, no significant difference was found in MKI67 and SOX2 expression between the latter three areas, thus suggesting the presence of a consistent tumor cell load also in the infiltrative zone.

To distinguish spots predominantly containing tumor cells from those comprising mostly stromal and immune cells, we inferred the copy number variation (CNV) profile from the transcriptome of each spot using the distal area as a reference. The vast majority of CNVs were observed in the core and inner edge of the tumor (**Figure S1G, Figure S1H**). The three patients share the gain of chromosome 7 (7amp) and the loss of chromosome 10 (10del) in the tumor samples and in a part of the spots of the outer edge (**Figure S1G, Figure S1H**). These are the most common CNVs of GBM. However, we also observed less common GBM CNVs that are patient-specific, such as the deletion of chromosome 9p in patient #1, and the deletion of chromosome 13 and 19 in patient #2^23,24^. We annotated spots containing either 7amp or 10del as cancer spots. Nearly all spots within the tumor core or inner edge were classified as cancer spots, whereas those at the outer edge displayed a mixture of cancer and normal spots. This pattern highlights the heterogeneous nature of the infiltrative zone (**Figure 1E**, **Figure 1F**), where cancer cells remain substantially represented, reaching up to 70% in patient #1.

After annotating spots as tumor and normal, we performed linear correlation of the tumor spots to investigate GBM regional heterogeneity. Interestingly, cancer spots in the core and inner edge of the tumor still cluster by patient, while tumor spots in the outer edge tend to cluster together, with higher reciprocal correlation (median Pearson correlation coefficient of 0.48 ± 0.11 for the outer edge, versus - 0.35 ± 0.18 and −0.09 ± 0.17 for the core and the inner edge, respectively) (**Figure 1G**).

This finding points toward a reduction in heterogeneity in the infiltrative region of GBM, suggesting a converging mechanism of invasion. The reduction in heterogeneity observed could be due either to a less diverse microenvironment, a selective advantage of a subpopulation of GBM cells in the infiltrative region, the need for specific mechanisms to invade the surrounding brain, or even a combination of all these factors.

### Inflammatory response is sharply reduced in the GBM infiltrative area

To elucidate the causes of the reduced heterogeneity of GBM infiltrative area, we next sought to disentangle the cell type composition of each spot of our spatial transcriptomics slides. Since one of the major limitations of the 10X Visium stRNA-seq platform is its inability to achieve single-cell resolution due to the size of the spatial gene expression spots, we analyzed the cell type composition of our normal and tumor spots by comparison with single cell profiles of GBM and healthy brain cells. We first generated a comprehensive reference single-cell dataset representative of both GBM cell states, GBM-associated non-tumor cells, and healthy brain cortex cells by integrating 38 scRNA-seq GBM samples from three different studies^3,4,7,8^ (**Figure 2A**). After filtering and batch-correction, clusters of transcriptionally similar cells were identified, projected onto a two-dimensional embedding and annotated based on established gene expression signatures of non-tumor cell types commonly found in GBM (**Figure S2A**). To better identify tumor cells, we inferred the CNV profile from the expression data (**Figure S2B**). Tumor cells were isolated, reclustered, and annotated using published gene signatures of GBM cells (**Figure S2C-E**)^3,4,7,8^. We identified one cluster with oligodendrocyte precursor-like cells (OPC-like) features, two clusters of neural progenitor-like cells (NPC-like), one cluster of astrocyte-like cells (AC-like), and two clusters of mesenchymal-like cells (MES-like), one with an hypoxia-dependent signature (MES1) and one hypoxia independent (MES1). Two clusters showed high expression of cell cycle markers and were therefore annotated as G1S and G2M cells (**Figure S2D**). We then integrated this GBM dataset with a pre-annotated, publicly available primary motor cortex single-nuclei RNA-seq dataset^25^ to obtain better representation of cell types commonly found in the healthy cortex (**Figure S2F, Figure 2A**).

**Fig. 2:**
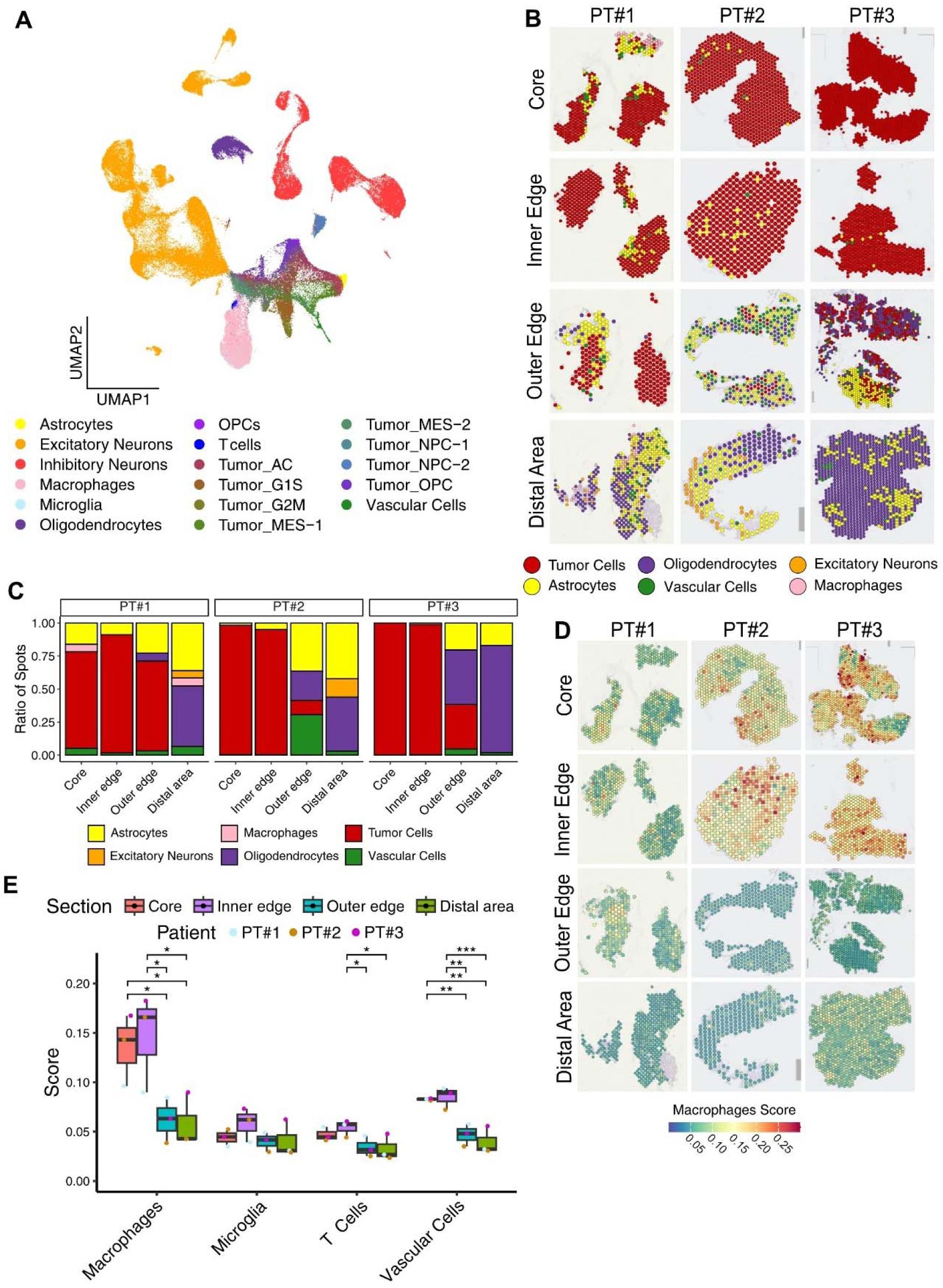
A gradient of inflammation from the core to the outer edge of GBMs. **(A)** Annotated UMAP of the integrated scRNA-seq reference dataset used to study spatial spot composition. The reference was obtained by integrating several published scRNA-seq GBM samples to an annotated public human motor cortex single-nuclei RNA-seq dataset. **(B)** Surface plots showing the predominant identity of spots previously assigned as normal. **(C)** Stacked barplots showing the ratio of each spot predominant identity over the total number of spots for each sample. **(D)** Surface plots of macrophage marker expression score in all spatial spots for all samples. **(E)** Boxplot with the average macrophage, microglia, T cell, and vascular cell scores in the four sampled areas. Dots represent the average score for each patient in that specific sampled area. One way ANOVA test with *post-hoc* Tuckey’s test. *p < 0.05, **p < 0.01, ***p < 0.001.

Exploiting our integrated reference of single-cell datasets, we first studied the features of our spatial transcriptomics spots annotated as normal. We projected the embeddings of the non-tumor spots onto the integrated reference latent space, revealing the predominant cell type comprising each spot (**Figure 2B**). The predominant non-tumor cell types in each area were glial cells, either astrocytes or oligodendrocytes, although in a limited number of spots vascular cells (endothelial cells and pericytes) and macrophages were the predominant population (**Figure 2C**). Neuron spots, exclusively of the excitatory type, were observed only in the distal area, likely due to the stressful environment inside and surrounding the tumor which makes neuron survival difficult (**Figure 2B**, **Figure 2C**).

Given that we detected few spots with predominant immune identity in both tumor and infiltrative areas, we sought to investigate the immune infiltration in our areas with a complementary, marker-based approach. In the core and inner edge of the tumor, we found higher expression of bone marrow-derived macrophages markers compared to microglia (**Figure 2D, Figure S2G**), which is consistent with previous findings^26,27^. The microglia score remained constant through our four areas, while we observed a significant decrease from core and inner edge to outer edge and infiltrative areas for the macrophages score (**Figure 2E**). This is likely a consequence of the difference in inflammation between bulk tumor areas and the infiltrative region, with macrophages preferentially infiltrating the tumor areas also due to increased vascularization. Indeed, we observed significantly higher levels of expression of vascular cell genes in the core and inner edge areas (**Figure 2E**) with niches dominated by vascular cell types (**Figure S2I**). Furthermore, the inner edge of the tumor had the highest score for T cells (**Figure 2E, Figure S2H**), suggesting that this area is the one where the immune system and the tumor interact the most.

This analysis highlights a gradient of inflammation and vascularization that peaks in the actively proliferating region of the tumor and then sharply decreases in the infiltrative area. The decrease in inflammation and the increase in glial cell types underlines a strong change in the microenvironment of infiltrative tumor cells, with expected impact on tumor invasiveness and relapse.

### GBM cells show pronounced oligodendrocyte lineage signatures in the infiltrative zone

We then exploited the integrated reference GBM scRNA-seq dataset to analyze the distribution of cancer cell states in our spatial dataset. By projecting tumor spots in our spatial samples on the low-dimensional space of the reference dataset, we observed a prevalence of AC-like and MES-1-like states in the core and inner edge of the tumor, whereas the OPC-like state was strongly enriched in cancer spots at the outer edge of the tumor (**Figure 3A**, **Figure 3B**, **Figure 3C**). We also observed the presence of niches of MES-2-like programs in the tumor core and, to a lesser extent, at the inner edge (**Figure S3A**). The MES-2-like cell state is generally associated with hypoxia, a prevalent feature of the tumor core, but in our samples this feature extends to the actively proliferating region of GBM. Finally, we noticed moderate prediction scores for the NPC-2-like state in a few spots in the core and inner edge of the tumor and spots of high prediction score in the outer edge of patient 2 (**Figure S3B**).

**Fig. 3:**
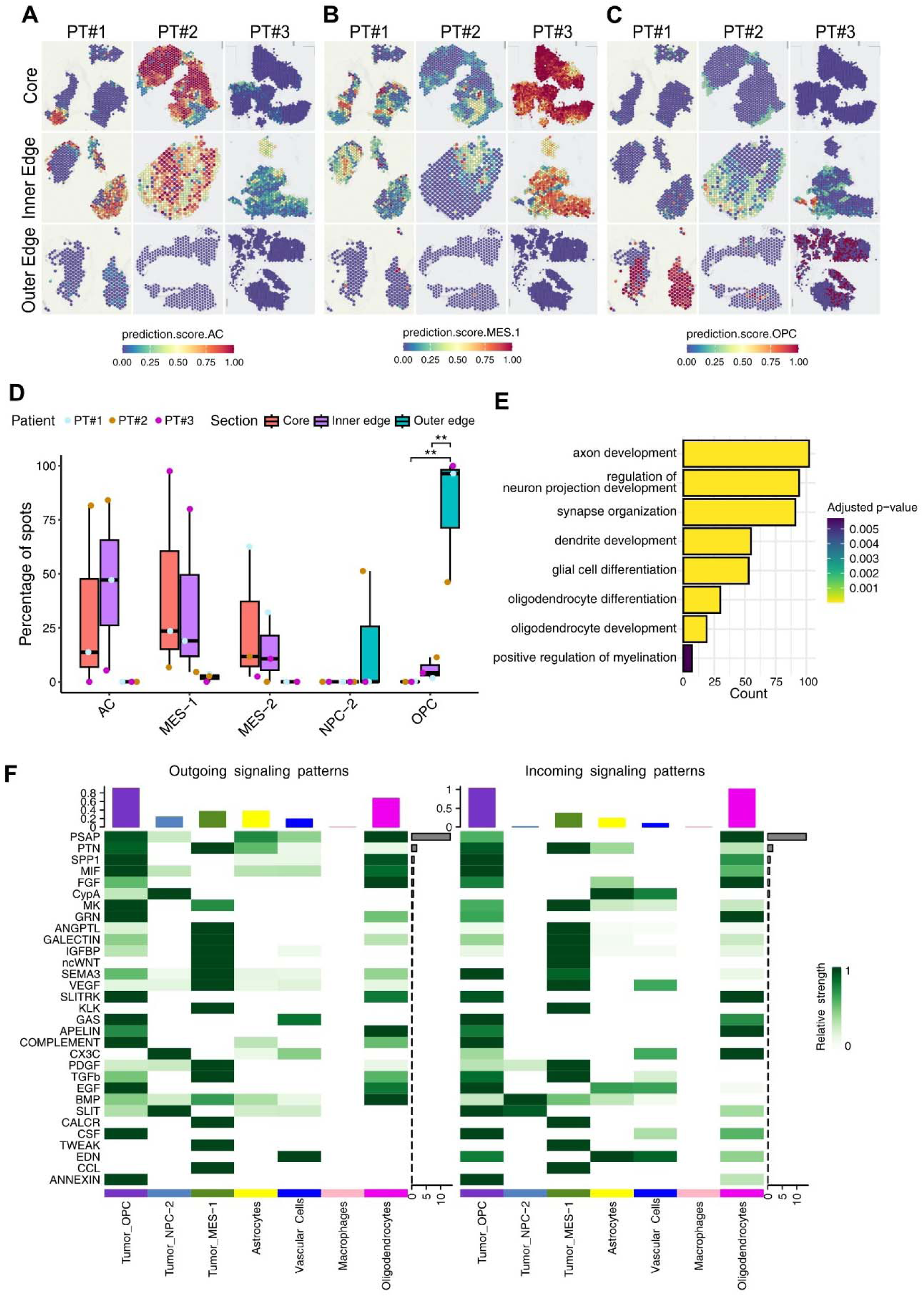
The OPC-like subpopulation is enriched in the outer edge of GBM. (A, B, C). Surface plots of AC **(A)**, MES1 **(B)**, and OPC **(C)** cell state prediction score in tumor spatial spots for all samples in the three tumor areas. The score represents the confidence of the predicted cell state on a scale from 0 to 1, based on the number of anchor genes found between the query spatial spot and the scRNA-seq reference embedding. **(D)** Boxplot showing the percentage of tumor spots mapped to each GBM cell state of the reference GBM scRNA-seq dataset. One way ANOVA test with post-hoc Tuckey’s test. **p < 0.01. **(E)** Barplot showing representative terms of the gene ontology enrichment analysis performed on genes differentially expressed by tumor spots in the outer edge of the tumor versus tumor spots in the other areas. **(F)** CellChat heatmaps showing the relative signaling strength of the most active signaling pathways in the different cell types of the outer edge of the tumor. The top histogram shows the total signaling strength of a cell type by summing the contribution of all of the pathways shown. The right histogram shows the total strength of a pathway by summing the strengths of all cell types shown. The left heatmap shows outward interactions, while the right heatmap shows inward interactions.

By assigning the identity of each tumor spot to the GBM state with the highest prediction score, we revealed a statistically significant increase of the OPC-like state within the cancer cell population in the outer edge compared to the other two tumor areas (**Figure 3D**). Indeed, tumor spots of the outer edge map very peripherally in the OPC-like state of the UMAP embedding, with patient #2 having also some spots mapping in the NPC2-like region (**Figure S3C, Figure S3D, Figure S3E**). Instead, spots of the core and inner edge of the tumor are much more colocalized, and spread between the AC-like and MES-like states (**Figure S3C, Figure S3D, Figure S3E**). These observations suggest that the shift of infiltrative GBM cells toward the OPC-like state is strong and cohesive, which is confirmed by the identity of genes overexpressed in outer edge tumor spots compared to tumor spots of the other areas (among which *SOX10*, *OPALIN*, *MYRF*, *MAL*, *NKX6-2*, *FA2H*) (**Figure S3F, Table S4**). The oligodendrocytic shift of tumor cells in the outer edge was further confirmed by gene over-representation analysis, revealing marked enrichment of neuronal projection and ensheathment programs (**Figure 3E, Table S4**). The same analysis for spots in the inner edge points to glial development, cell adhesion and immune modulation (**Figure S3G, Table S4**), while spots in the core of the tumor are instead enriched for metabolic signatures, as well as terms of response to cell hypoxia and stress (**Figure S3H, Table S4)**. The oligodendrocyte signature is therefore a predominant feature of tumor cells of the tumor infiltrative zone. To understand if the observed shift toward the OPC-like state in GBM infiltrative region might be a consequence of the different microenvironment to which tumor cells are exposed, we sought to infer signaling involved in the infiltrative zone of GBM. We predicted ligand-receptor interactions from the transcriptomes of our spots in the inner edge and outer edge of the tumor, using both spot identities assigned in our tumor and non-tumor analyses. Overall, ligand-receptor signaling in both the inner edge and outer edge of the tumor is heavily tilted in favor of the most abundant spot types, namely AC-like and MES-2-like for the former, and OPC-like for the latter (**Figure S3I**). However, we observed stronger interactions between tumor and nontumor spots in the outer edge, in particular from and toward astrocytes, vascular cells, and normal oligodendrocytes. In the inner edge of the tumor, the strongest signaling pathways mostly involve immunomodulatory and pro-survival cytokines, such as PTN, SPP1, MK, and CypA, and also mitogenic signaling such as FGF, EGF, and VEGF (**Figure S3J, Table S5**). Interestingly, PTN and its ligand PTRPZ1 have been linked to radial glia-like behavior in GBM cells^6^. Instead, the signaling network of the outer edge is dominated by the prosaposin (PSAP) pathway (**Figure 3F**), mostly driven by the interaction between the PSAP ligand and the GPR37 receptor (**Table S5**). In the outer edge, PSAP signaling is largely happening between normal oligodendrocyte spots, as marked by the high interaction probability, but strong interactions also occur between tumor OPC-like spots and normal oligodendrocyte spots. From one side, the PSAP pathway has been reported to have a broad effect on lipid metabolism, proliferation, survival, and immunomodulation and to have a neuroprotective role in the nervous system^28^. However, from the other side, it has also been implicated in cancer progression in the context of metastasis^29^ and in oligodendrocyte-driven neuroinflammation in neurodegenerative disorders^30^. This signaling might constitute a selective advantage of the OPC-like cancer population that allows it to thrive in the infiltrative areas, further invade the surrounding brain, and later give rise to recurrence.

Overall, these data elucidate regional differences in GBM cell states, with cells in the necrotic core and proliferative edge acquiring MES-like and AC-like phenotype, while cells in the infiltrative margin adopting a OPC-like state, with consequently altered interaction with neighboring cells, both tumor and non-tumor.

### The Prevalent Oligodendrocytic Profile of Infiltrating GBM Cells is a General Feature

To further validate our observation of the OPC-like features of the infiltrative zone and to exclude that our results might be due to patient-specific characteristics combined with limited sample size, we decided to corroborate these data using complementary approaches (**Figure 4A**).

**Fig. 4:**
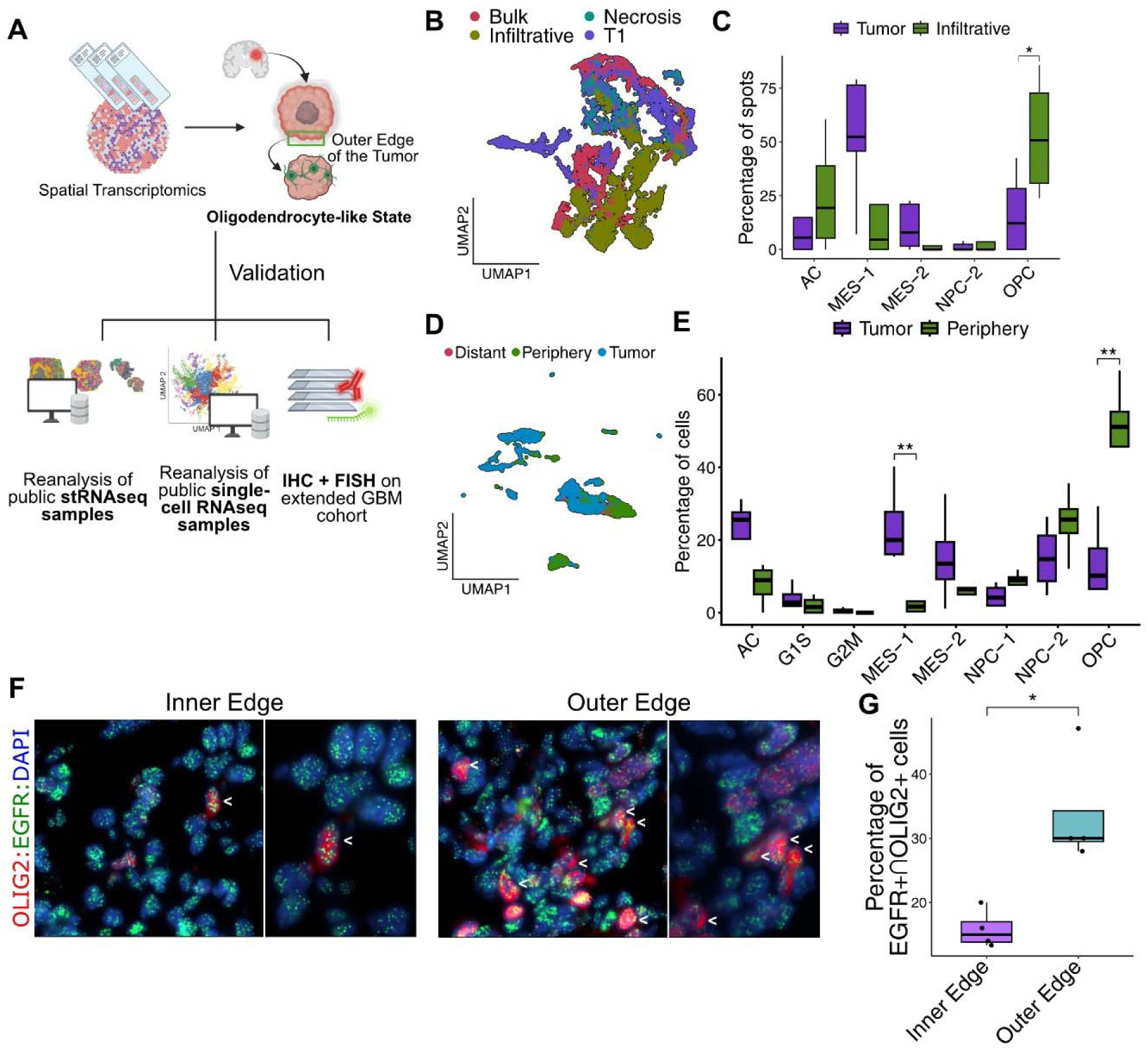
The OPC feature of infiltrative GBM cells is prevalent in two distinct stRNA-seq and scRNA-seq datasets and in IHC-FISH samples. **(A)** Scheme of the analyses performed to validate the predominant OPC-like features of GBM cells in the infiltrative zone. **(B)** UMAP plot of all integrated spots from the Greenwald et al. spatial transcriptomics dataset. Spots are colored by sampled area. **(C)** Boxplot showing the percentage of spots of the Greenwald et al. dataset from the tumor areas (including the bulk, T1, and necrosis areas) or the infiltrative area mapped to the GBM scRNA-seq dataset. One way ANOVA test with post-hoc Tuckey’s test. *p < 0.05. **(D)** UMAP of all integrated cells from the Darmanis et al. scRNA-seq dataset. Spots are colored by sampled area. **(E)** Boxplot showing the percentage of cells of the Darmanis et al. dataset from the tumor or the infiltrative area mapped to the GBM scRNA-seq dataset. One way ANOVA test with post-hoc Tuckey’s test. **p < 0.01. **(F)** Representative images of IHC + FISH for OLIG2 and *EGFR* on tissue slides from the inner edge and the outer edge of the tumor. The arrows indicate cells of high OLIG2 expression and *EGFR* gene amplification. **(G)** Boxplot showing the percentage of *EGFR* amplified and OLIG2 positive cells for all samples analyzed of the inner edge and outer edge of the tumor. Two-sided t-test. *p < 0.05.

First of all, we reproduced our analysis using a recently published spatial transcriptomics dataset^17^, comprising GBM samples of comparable numerosity to ours. In detail, this dataset includes six samples from the tumor bulk, one sample from the necrotic region (corresponding to our core), four samples from the T1 region (roughly our inner edge), and four samples from the infiltrative zone (**Figure 4B, Figure S4A, Figure S4B, Figure S4C**). We scored cell spots with the signatures derived from our integrated GBM scRNA-seq dataset and found that mesenchymal and astrocytic clusters characterized the bulk, T1, and necrotic areas, whereas the infiltrative zone showed a significant enrichment of OPC-like spots, consistent with our findings (**Figure 4C**).

Next, we validated our results using an external scRNA-seq dataset comprising four GBM patients^5^. Each patient included one sample from the tumor bulk, roughly corresponding to our core and inner edge, and one sample from the peritumoral brain, corresponding to our outer edge (**Figure 4D, Figure S4D, Figure S4E, Figure S4F**). As for the spatial transcriptomics datasets, we projected the embeddings of these scRNA-seq datasets onto the low-dimensional space of our GBM reference. The OPC-like cluster was predominantly observed in the infiltrative zone, located at the periphery of the tumor (**Figure 4E**), further confirming our previous findings. Moreover, in this dataset, the tumor region exhibited a predominance of AC-like and MES-like subpopulations, along with the presence of the OPC-like cluster (**Figure 4E**).

Finally, we confirmed the increased OPC-like features of infiltrative GBM cells both *via* IHC on tissue sections near-adjacent to those used in our stRNA-seq analysis and via combined IHC and fluorescence *in situ* hybridization (FISH) analysis on samples derived from additional 5 GBM patients. IHC on tissue sections near-adjacent to those used in our stRNA-seq analysis revealed increased OLIG2 expression in the outer edge of the tumor compared to the core and the inner edge (**Figure S4G**, **Figure S4H**). FISH analysis of the *EGFR* gene combined with IHC for the OLIG2 protein enabled us to distinguish between OLIG2 signals originating from tumor cells and those originating from non-tumor OPCs and oligodendrocytes. Specifically, among GBM cells identified as *EGFR* amplified (EGFR+) cells, we counted those positive for OLIG2 in samples both from the inner and outer edges of the tumor (**Figure 4F**). In the inner edge, we observed an average of 16 ± 3% of OLIG2+ cells among all cells with *EGFR* amplification (**Figure 4F**, **Figure 4G**). Of note, in this tumor area, all OLIG2+ cells, albeit representing a minority of the neoplastic bulk, carried *EGFR* CNVs, suggesting that normal OPCs are absent or barely detected in the inner edge. Conversely, in the outer edge, we detected an average of 34 ± 9% OLIG2+ GBM cells (**Figure 4F**, **Figure 4G**), indicating an increasing density of GBM cells with an OPC-like profile. Contrary to what we observed in the inner edge, the EGFR+/OLIG2+ cells identified in the outer edge represented a minority of all OLIG2+ cells, suggesting the presence of a mixed cell population mainly composed of normal appearing oligodendrocytes and OPCs.

In summary, by using data from both scRNA-seq and stRNA-seq datasets, as well as IHC and FISH-IHC on our samples, we proved that the enrichment of the OPC-like cluster in the infiltrative zones of GBM is a general feature. The reported phenotypic shift of infiltrative GBM cells represents a therapeutically exploitable vulnerability that opens the possibility of targeting the OPC-like state of GBM cells to prevent tumor relapse, which remains the leading cause of patient death.

## Discussion

GBM remains one of the most lethal tumors, with nearly inevitable relapse even after aggressive therapeutic treatments. GBM is characterized by strong heterogeneity, both within different regions of the same tumor and between patients. This spatial and inter-patient heterogeneity poses a major challenge to treatment. In particular, the infiltrative tumor margin, where residual cells often evade surgical resection and initiate recurrence, represents a critical yet understudied area. Thorough regional sampling and molecular characterization are therefore essential to better understand the cellular states driving tumor progression and relapse.

In this context, we performed spatial transcriptomics on four surgically defined tumor regions: the tumor “core” (central necrotic area), the “inner edge” (characterized by strong Gliolan fluorescence), the “outer edge” (the infiltrative zone, defined by low Gliolan fluorescence and FLAIR hyperintensity), and the “distal area” (healthy tissue distant from the tumor). We found that core and inner edge regions from the same patient were more similar to each other than to corresponding regions in other patients, highlighting the well-known high inter-patient heterogeneity of GBM. In contrast, malignant spots from the outer edge exhibited a greater degree of similarity across patients, suggesting a conserved transcriptional program in infiltrating cells.

To decipher the cellular composition of each region, we integrated our spatial data with a single-cell reference atlas of GBM and healthy cortex. Interestingly, cells located in the central tumor regions (i.e. core and inner edge) were enriched for astrocyte-like (AC-like) and mesenchymal (MES-like) states, consistent with previous findings associating these cell states with hypoxic stress and therapy resistance^3,8,17,31^. GO enrichment revealed upregulation of hypoxia response, inflammatory, and metabolic adaptation pathways. These results reinforce the concept that the tumor core harbors cells adapted to environmental stress through activation of compensatory pathways, including oxidative metabolism and pro-survival signaling. The inflammatory environment of the core and inner edge, shaped by autocrine and paracrine signaling as well as by interactions with immune cells, likely contributes to the dominance of MES- and AC-like states in the tumor bulk areas. Indeed, prior studies have shown that inflammation and immune cell interactions can promote proneural-to-mesenchymal transitions in GBM, enhancing resistance and invasiveness^32–34^.

While much focus has traditionally been placed on the tumor bulk, increasing attention has been recently directed toward the GBM infiltrative area both from a biological and clinical perspective^5,21,22^. Indeed, the unparalleled tumor regenerating ability of GBM, which causes recurrence in almost all patients^35^, is mostly due to invasive cells from the resection margin that not only escape surgery but also resist chemoradiation^36^. As these cells are the ultimate cause of the deadly prognosis of GBM, studying their degree of stemness, plasticity and invasivity, as well as the cellular ecosystem in which they regenerate the tumor is critical for developing more effective therapies^37^.

From our analysis, tumor cells present in the tumor infiltrative area (i. e. outer edge) exhibit oligodendrocyte progenitor cell-like (OPC-like) expression profiles, supported by enrichment for genes involved in oligodendrocyte lineage development. This finding is further strengthened by validation using external datasets, i.e. the spatial transcriptomics dataset from Greenwald and the scRNA-seq dataset from Darmanis^5,17^, both confirming a high abundance of OPC-like cells in the outer edge of GBM. OPC-like cells are a well-characterized population in GBM and are known for their marked plasticity and ability to adapt to various microenvironments^3,6,8^.

The molecular mechanisms driving the shift of GBM cells toward an OPC-like phenotype remain largely unclear. To explore potential microenvironmental influences, we analyzed tumor–microenvironment interactions at the outer edge and observed increased proximity between OPC-like GBM cells and surrounding oligodendrocytes, astrocytes, and vascular cells. We hypothesize that signaling molecules released by these neighboring cells may contribute to the proneural shift of infiltrating tumor cells. Supporting this idea, prior work has suggested that the white matter niche promotes OPC-like differentiation in infiltrating GBM cells, possibly as a response to injury^38^. In that study, SOX10 was proposed as a key regulator of this lineage shift. Here, we extend these findings by implicating PSAP signaling as an additional factor. PSAP is known to be upregulated in response to nerve injury, ischemia, or compression^39^, and has been shown to promote invasion in GBM in vitro. Although previously associated with the mesenchymal GBM subtype^40^, our data suggest that PSAP may also play a role in the invasive behavior of OPC-like cells. We propose that tumor-induced mass effect and consequent brain edema may provoke an injury response in the surrounding tissue, leading to increased release of stress signals such as PSAP from non-tumor cells. GBM cells infiltrating these areas may then be exposed to PSAP, which could enhance their migratory capacity and promote survival. However, to date, no direct link has been established between PSAP signaling and the induction of an OPC-like transcriptional program. Thus, the mechanisms underlying the oligodendrocytic shift of GBM cells in this context remain to be fully elucidated.

A key question emerging from our study is how infiltrative OPC-like GBM cells transition into other cell states to initiate tumor relapse. Recent single-cell analyses of matched primary and recurrent GBM samples post-therapy^31,41^ have begun to address this issue. One such study reports a shift in cell state composition, primarily toward a mesenchymal phenotype, implying that residual cells escaping resection and evading therapy retain the capacity to reconstruct the original tumor architecture. In this context, radiotherapy and chemotherapy may contribute to an inflammatory microenvironment that promotes the reprogramming of OPC-like GBM cells into mesenchymal or astrocyte-like states^42–44^.

Overall, the mechanisms driving invasion and cell state plasticity in GBM are complex, involving dynamic interactions among malignant cells, therapeutic interventions, and the surrounding brain microenvironment. Future work should prioritize the development of experimental models that faithfully recapitulate the tumor-adjacent brain niche to better dissect its role in shaping GBM cell behavior.

In summary, our study reveals region-specific cellular states in GBM, with a striking conservation of OPC-like cells at the infiltrative margin. These cells, through their plasticity and positioning, may be central drivers of recurrence. Targeting their differentiation pathways or exploiting their vulnerabilities could represent a promising therapeutic avenue. Future efforts should aim to recreate the invasive niche in experimental models, enabling functional studies of the microenvironmental signals that shape GBM cell states and behaviors.

## Methods

### Patient samples

Tumor samples were obtained from patients with GBM who had undergone surgical resection at Santa Chiara Hospital in Trento. The local ethics committee of the Santa Chiara Hospital in Trento approved the use of patient samples for this study. All enrolled patients were fully informed and signed the preoperative informed consent form. Clinical characteristics are summarized in **Table S1**.

Histopathological analyses of tumor sections, summarized in **Table S1**, were performed by the Pathology Department of Santa Chiara Hospital. Experienced neuropathologists reviewed tissue sections and tumor types were diagnosed according to the World Health Organization 2021 classification of central nervous system tumors^46^.

### Tissue collection, handling, and Visium spatial gene expression protocol

Immediately after resection, tissue samples were snap-frozen in isopentane pre-cooled with liquid nitrogen. Frozen samples were immediately transported to the laboratory and stored at −80°C. Frozen samples were embedded in OCT and stored at −80°C until further processing.

For each area, 10-15 sections were pooled and total RNA was extracted using Trizol. RNA quality was assessed using the Agilent 2100 Bioanalyzer according to the manufacturer’s instructions. The correct permeabilization time for the Visium Spatial Gene Expression protocol was determined for each area with the 10X Visium spatial tissue optimization kit (PN-1000193).

Spatial gene expression experiments were performed using the fresh-frozen 10X Visium Spatial Gene Expression Kit (PN-1000184). Briefly, tissues embedded in OCT were sectioned on a cryostat at −20°C. 10 μm thick tissue sections were placed on the Spatial Gene Expression slides. One tissue section per area was placed on a Visum Spatial Gene Expression capture area. Specifically, the four sampled areas from each patient were placed on the four capture areas of the same slide. Visium slides were fixed in pre-cooled 100% methanol and H&E staining was performed according to the 10X Genomics protocol. Mosaic brightfield images were captured using a Zeiss Axio Imager M2 microscope at 10x magnification.

The brightfield configuration includes the use of a color camera (3×8 bit, 2424×2424 pixel resolution), white balance functionality, 2.18 µm/pixel capture resolution, and 2-10 millisecond exposure times. Permeabilization was performed according to the manufacturer’s instructions using the predefined time for each area as determined by the Tissue Optimization Protocol. RNA was then reverse transcribed, a second strand was produced and cDNA was amplified using the appropriate number of PCR cycles as determined by qPCR. The cDNA quality was evaluated using an Agilent Bioanalyzer high-sensitivity chip. cDNA was amplified for 18 cycles and purified using AMPure beads. Enzymatic fragmentation and two-sided size selection were performed. P5, P7, i7, and i5 samples and TruSeq Read 2 were added by end repair, A-tailing, adapter ligation, and PCR. AMPure beads were used for post-ligation clean-up. P5 and P7 indexes were added by PCR and the final library was size selected using AMPure beads. The libraries were sequenced on an Illumina NovaSeq 6000 using the Novaseq S1 100 cycles sequencing kit (Illumina, Cat#20028319).

### Immunohistochemistry on GBM tissue sections

Tissue samples obtained from the different tumor areas were fixed in 4% paraformaldehyde, cryoprotected in 30% sucrose in PBS, snap frozen in liquid nitrogen and 5μm cryostat sections were obtained. Tissue sections taken from a −20°C freezer were left at room temperature for 30 minutes and then washed in PBS. The sections were permeabilised using 1% SDS in 0.3% Triton X-100 in PBS for 20 minutes. After a wash in PBS, the tissue samples were incubated overnight at 4°C with primary antibodies diluted in a blocking solution containing 3% goat serum and 0.3% Triton X-100 in PBS. The concentration of each primary antibody is specific, OLIG2 1:500 (Genetex GT132732), SOX2 1:200 (Santa Cruz SC365823), MKI67 1:500 (Abcam AB15580). The next day, after a wash in PBS, the slides were incubated with AlexaFluor secondary antibodies diluted 1:500 for 1.5 hours at room temperature (anti-mouse for SOX2, Thermo Fisher Scientific Cat#A11070; anti-rabbit for OLIG2 and MKI67, Thermo Fisher Scientific Cat#A21442). The nuclei were stained using Hoechst 33342 (1 µg/ml, Thermo Fisher Scientific, Cat#H1399). Coverslips were applied to the slides, which were then stored at +4°C until imaging. Slide acquisition was performed using an SP8 Leica confocal microscope. Immunopositive cells were quantified by counting positive nuclei and normalising to the number of Hoechst-positive nuclei.

### Fluorescence in situ hybridization combined with immunofluorescence on GBM tissue sections

Tumor samples from five GBM patients were analyzed by FISH for *EGFR* copy number detection combined with IHC for OLIG2 expression. IHC analysis for OLIG2 was performed using a rabbit anti-OLIG2 Rabbit monoclonal primary antibody 1:600 (Roche Diagnostics, Cat#07667973001). After washing, cells were incubated with an Alexa Fluor® 488 anti-rabbit secondary antibody (Thermo Fisher Scientific Cat#A11094). Fluorescent images were captured using the Image-Pro Plus software. After IHC evaluation, the same tissue sections were used to perform combined fluorescence in situ hybridization (FISH) for *EGFR* copy number detection. FISH was performed according to standard protocols using the Zytolight SPEC EGFR/CEN7 Dual Color Probe fluorescent probe (Zytovision GmbH, Bremerhaven, Germany), which is composed of ZyGreen labeled polynucleotides (excitation 503 nm/emission 528 nm), targeting sequences in 7p11.2 harboring the *EGFR* gene region, and ZyOrange labeled polynucleotides (excitation 547 nm/emission 572 nm), targeting sequences mapping in 7p11.1-q11.1 specific for the alpha satellite centromeric region D7Z1 of chromosome 7. The sections were then counterstained with DAPI/Antifade Solution. The slides were evaluated by an expert cytogeneticist using a Zeiss Axioscope system (Zeiss, Milan, Italy), and a minimum of 50 tumor nuclei for each sample were analysed by the Metafer v4.1.1 software, which is able to automatically count probe signals (MetaSystems s.r.l., Milan, Italy). *EGFR* positive cells were identified as cells with ZyGreen cluster amplification, defined as a highly uncountable increase of signals in ≥5% of analyzed cells. CEN7 copies were not counted as CEN7 ZyOrange probe signals were obscured by Alexa Fluor® 488 IHC OLIG2 expression. OLIG2 positive or negative tumor cells were identified as cells carrying *EGFR* copy number alterations, while OLIG2 positive cells without any *EGFR* alterations have been considered as non-neoplastic oligodendrocytes.

### Spatial data preprocessing and tumor spot identification

Raw sequencing reads from spatial experiments were processed using the standard pipeline for fresh frozen tissues provided by Space Ranger software v2.0.1 specifically developed for Visium Spatial gene Expression data processing. Alignment was performed using human transcriptome reference GRCh38. The obtained gene expression matrices were analyzed with the R package Seurat v4.4.0 . Sample quality controls included the evaluation of UMI and gene count together with the cell mitochondrial read content. Low quality spots with number of reads smaller than 100, number of unique expressed genes smaller than 100, and mitochondrial gene ratio larger than 0.40 were excluded from downstream analysis.

Spatial spot gene expression was normalized using the SCTranform Seurat pipeline. Samples were integrated using Canonical Correlation Analysis (CCA). We ran PCA analysis and UMAP projection of the integrated samples selecting the first 15 principal components. NMF dimensional reduction analysis was performed using the R package Singlet with the function RunNMF() while we used RunLNMF() function on the merged spatial object to account for batch effects. To evaluate similarity between the different tissue sections we computed Pearson’s correlation on each sample normalized gene expression data using the do_CorrelationPlot() function of the SCpubr v2.0.2 R package^48^. We used the inferCNV R package^49^ to study the copy number alteration profile of the tissue areas. We ran the analysis separately for each patient to mitigate any possible batch effects in the analysis. For each patient, the distal area sample was used as reference. InferCNV was run with the following parameters: analysis_mode=“subclusters”, cutoff=0.1, denoise=TRUE, HMM=F and k_obs_groups=1. Gene positions were taken from Gencode human assembly release 44 (GRCh38.p14). A chromosome region was defined as amplified or deleted if the associated inferCNV score for that specific region was smaller or greater than 1, respectively. We defined tumor spots as those harboring the amplification of chromosome 7 or the deletion of chromosome 10.

### Generation of an integrated scRNA-seq dataset

Single-cell RNA-seq datasets of IDH wild type glioblastoma samples from three published studies^3,8,50^ were retrieved from the Gene Expression Omnibus database. To ensure consistency in the analyses, we retained samples processed with the 10X Genomics platform, resulting in a total of 38 GBM samples. Downstream computational analyses were conducted using the Seurat v4.4.0 R package. For GBM samples, we removed low-quality cells with fewer than 800 UMIs, fewer than 300 detected genes, or a mitochondrial UMI content exceeding 30% of total UMI counts. Cell doublets were predicted using the DoubletFinder v2.0.3 R package^51^. Samples with less than 1000 cells after filtering were excluded from downstream analysis, resulting in 27 samples comprising approximately 69,000 cells, and data were then processed using the standard Seurat pipeline with default parameters. The cell cycle phase was predicted with the CellCycleScoring() function in Seurat. GBM samples were integrated via CCA to remove the batch effect. Transcriptionally similar clusters of cells were then annotated using gene expression scores for markers of nontumor cells commonly found in GBM, namely innate immune cells (*AIF1*, *FCER1G*, *TYROBP*, *ITGAM*) oligodendrocytes (*PLP1*, *MBP*, *MAG*, *MOG*), T cells (*PTPRC*, *CD3E*, *CD4*, *CD8A*, *CD2*, *CD3A*, *CD3G*), pericytes (*ACTA2*, *PDGFRB*), and endothelial cells (*PECAM1*, *VWF*, *APOLD1*, *CLDN5*). Macrophages and microglia cells were further annotated via subclustering and expression of more specific markers, in particular *TMEM119*, *CX3CR1*, *TREM2* for macrophages, and *CD14*, *CD163*, *LYZ*, *S100A8* for microglia. Gene expression scores were computed using the AddModuleScore_UCell function of the UCell v3.21 R package^52^.

CNV inference was performed separately for each sample via the inferCNV package using a subset of cells in the macrophage cluster with high expression of macrophage markers as reference. We considered a chromosome region amplified or deleted if the associated inferCNV score for that specific region was larger than 1.05 or smaller than 0.95, respectively. We defined tumor cells by combining the results from the CNV analysis, clustering, and lack of non-tumor marker expression.

Isolated tumor cells were then re-integrated via reciprocal PCA and re-clustered with a resolution parameter of 0.8. Cell clusters were then annotated using the scType v1.0 R package^53^ by using gene lists for GBM states published in Neftel et al. 2019^3^.

Pre-filtered and pre-annotated human M1 primary motor cortex 10X Genomics scRNA-seq data were downloaded from the Allen Brain Atlas. Data were processed using the standard Seurat pipeline and were then integrated with GBM datasets using reciprocal PCA.

### Spatial spot gene expression scoring

Spot gene expression scores for nontumor cell types were computed using the AddModuleScore_UCell() function of the UCell v3.21 R package using the top 80 differentially expressed genes ordered by log_2_ fold change of nontumor clusters in the reference scRNA-seq dataset. Differentially expressed genes were determined via the FindAllMarkers() function of Seurat with default parameters. Both spots annotated as tumor and nontumor were scored using this method.

### Projection analysis

Projection of spatial transcriptomic spots onto the reference scRNA-seq dataset was performed using the FindTransferAnchors() and MapQuery() functions in Seurat with default parameters, projecting the low-dimensional space of each sample onto the reciprocal PCA embedding of the reference dataset. Projection of spots annotated as normal was performed with nontumor cell types in the reference, while spots annotated as tumor were projected onto tumor cell states of the reference dataset. The same method was used for the projection of the Greenwald et al. and the Darmanis et al. datasets.

### Differential expression and gene ontology enrichment analyses

We performed differential expression analyses between spatial spots using the Seurat function FindAllMarkers() with default parameters. Gene Ontology enrichment was performed with the ClusterProfileR v4.6.2 R package^54^.

### Cell-Cell Communication Analysis

Ligand-receptor interaction analysis among spatial spots was performed using the CellChat v2.2.0 R package^54^. In detail, we created a CellChat object for each of the analyzed areas, using the three patients as replicates and we inferred ligand-receptor interactions using the built-in database subset for secreted signaling. We then defined the overexpressed interactions with the identifyOverExpressedGenes() and identifyOverExpressedInteractions() functions and determined ligand-receptor interactions via CellChat probabilistic approach with the computeCommunProb() function, using parameters type = “truncatedMean”, trim = 0.1, distance.use = TRUE, interaction.range = 250, scale.distance = 0.01, contact.dependent = TRUE, contact.range = 100. Interactions supported by less than 5 spots were filtered out. Interactions were then summarized at the pathway levels using the computeCommunProbPathway() function and aggregated with the aggregateNet() function. Results were visualized using the netVisual_circle() and the netAnalysis_signalingRole_heatmap() functions. CellChat analyses were performed separately for either the inner edge or outer edge of the tumor samples.

### Statistical information

Data analysis was conducted using R v4.3.0. No statistical method was used to predetermine sample size. For multiple group comparisons, analysis of variance (ANOVA) using the F test was employed to compare the two groups and p-values were computed *via post-hoc* Tukey’s test. We applied parametric tests, including Student’s t-test (two-tailed). When parametric tests were used, we did not formally test for normality. Statistical significance was set at p-value[<[0.05.

## Data Availability

The spatial transcriptomics RNA sequencing raw data generated in this study, along with counts matrices and metadata for each sample, are publicly available in GEO under accession code GSE309418. The Neftel et al. dataset and Wang et al. datasets were downloaded from GEO using accession codes GSE131928 and GSE138794, respectively. The Couturier et al. dataset was obtained directly from the authors upon request. The human primary motor cortex dataset was obtained from the Allen Brain Atlas (https://portal.brain-map.org/atlases-and-data/rnaseq/human-m1-10x).

The Greenwald et al. and the Darmanis et al. were obtained from GEO with the accession codes GSE237183 and GSE84465, respectively.

## Code Availability

No custom computer code was used for the analyses included in this article.

## Contributions

A.Q. designed and supervised the study. C.M.A. performed tissue processing; C.M.A., D.S., S.L. and

A.S. performed library preparations; C.M.A. performed in vitro assays; under the supervision of A.Q.

Glioblastoma samples used in this study were acquired by C.M.A at the “S. Chiara” Hospital of Trento, Italy, under the supervision of S.S., F.C., and L.A.

S.S. and F.C., and L.A. identified cases for inclusion in the study. M.B. performed the pathological assessment on surgical samples.

E.S. and E.F.R. performed all bioinformatics analyses under the supervision of A.Q. and T.T.

L.P. performed the in-situ fluorescence hybridization experiments and image analysis under the supervision of P.L.P. A.Q., C.M.A., E.S., E.F.R., and D.S. wrote the manuscript with input from all authors. All authors read and approved the final manuscript. Corresponding author Correspondence to: Author-4

Corresponding authors must provide their ORCID ID before resubmitting the final version of the manuscript. Non-corresponding authors are encouraged to link their ORCID.

## Acknowledgements (optional)

The authors thank the Next Generation Sequencing, the Cell Analysis and Separation, and the Imaging Core Facilities of the Department of Cellular, Computational and Integrative Biology (CIBIO) for assistance with experimentation.

## Ethics declarations

### Competing interests

The authors declare no competing interests.

## Notes

### Competing Interest Statement

The authors have declared no competing interest.

## References

1. Kanderi, T. & Gupta, V., Glioblastoma Multiforme. in StatPearls (StatPearls Publishing, Treasure Island (FL), 2022).

2. Bette, S. et al. Retrospective analysis of radiological recurrence patterns in glioblastoma, their prognostic value and association to postoperative infarct volume. Sci. Rep. 8, 4561 (2018).

3. Neftel, C. et al. An Integrative Model of Cellular States, Plasticity, and Genetics for Glioblastoma. Cell 178, 835–849.e21 (2019).

4. Richards, L. M. et al. Gradient of Developmental and Injury Response transcriptional states defines functional vulnerabilities underpinning glioblastoma heterogeneity. Nat Cancer 2, 157–173 (2021).

5. Darmanis, S. et al. Single-Cell RNA-Seq Analysis of Infiltrating Neoplastic Cells at the Migrating Front of Human Glioblastoma. Cell Rep. 21, 1399–1410 (2017).

6. Bhaduri, A. et al. Outer Radial Glia-like Cancer Stem Cells Contribute to Heterogeneity of Glioblastoma. Cell Stem Cell 26, 48–63.e6 (2020).

7. Wang, R. et al. Adult Human Glioblastomas Harbor Radial Glia-like Cells. Stem Cell Reports 15, 275–277 (2020).

8. Couturier, C. P. et al. Single-cell RNA-seq reveals that glioblastoma recapitulates a normal neurodevelopmental hierarchy. Nat. Commun. 11, 3406 (2020).

9. Venteicher, A. S. et al. Decoupling genetics, lineages, and microenvironment in IDH-mutant gliomas by single-cell RNA-seq. Science 355, eaai8478 (2017).

10. Yabo, Y. A., Niclou, S. P. & Golebiewska, A. Cancer cell heterogeneity and plasticity: A paradigm shift in glioblastoma. Neuro. Oncol. 24, 669–682 (2022).

11. Sun, W. et al. Spatial transcriptomics reveal neuron-astrocyte synergy in long-term memory. Nature 627, 374–381 (2024).

12. Ståhl, P. L. et al. Visualization and analysis of gene expression in tissue sections by spatial transcriptomics. Science 353, 78–82 (2016).

13. Ravi, V. M. et al. T-cell dysfunction in the glioblastoma microenvironment is mediated by myeloid cells releasing interleukin-10. Nat. Commun. 13, 925 (2022).

14. Ravi, V. M. et al. Spatially resolved multi-omics deciphers bidirectional tumor-host interdependence in glioblastoma. Cancer Cell 40, 639–655.e13 (2022).

15. Ren, Y. et al. Spatial transcriptomics reveals niche-specific enrichment and vulnerabilities of radial glial stem-like cells in malignant gliomas. Nat. Commun. 14, 1028 (2023).

16. Zheng, Y., Carrillo-Perez, F., Pizurica, M., Heiland, D. H. & Gevaert, O. Spatial cellular architecture predicts prognosis in glioblastoma. Nat. Commun. 14, 4122 (2023).

17. Greenwald, A. C. et al. Integrative spatial analysis reveals a multi-layered organization of glioblastoma. Cell 187, 2485–2501.e26 (2024).

18. Lv, X. et al. Decoding heterogeneous and coordinated tissue architecture in glioblastoma using spatial transcriptomics. iScience 27, 110064 (2024).

19. Haddad, A. F., Young, J. S., Morshed, R. A. & Berger, M. S. FLAIRectomy: Resecting beyond the contrast margin for glioblastoma. Brain Sci. 12, 544 (2022).

20. Weller, M. et al. EANO guidelines on the diagnosis and treatment of diffuse gliomas of adulthood. Nat. Rev. Clin. Oncol. 18, 170–186 (2021).

21. Zigiotto, L. et al. Maximizing tumor resection and managing cognitive attentional outcomes: Measures of impact of awake surgery in glioma treatment. Neurosurgery (2025) doi:10.1227/neu.0000000000003591.

22. Niyazi, M. et al. ESTRO-EANO guideline on target delineation and radiotherapy details for glioblastoma. Radiother. Oncol. 184, 109663 (2023).

23. McNulty, S. N. et al. Beyond sequence variation: assessment of copy number variation in adult glioblastoma through targeted tumor somatic profiling. Hum. Pathol. 86, 170–181 (2019).

24. Mirchia, K. et al. Total copy number variation as a prognostic factor in adult astrocytoma subtypes. Acta Neuropathol Commun 7, 92 (2019).

25. Bakken, T. E. et al. Comparative cellular analysis of motor cortex in human, marmoset and mouse. Nature 598, 111–119 (2021).

26. Chen, Z. et al. Cellular and molecular identity of tumor-associated macrophages in glioblastoma. Cancer Res. 77, 2266–2278 (2017).

27. Musca, B. et al. The immune cell landscape of glioblastoma patients highlights a myeloid-enriched and immune suppressed microenvironment compared to metastatic brain tumors. Front. Immunol. 14, 1236824 (2023).

28. Yan, L. et al. Dissecting the roles of prosaposin as an emerging therapeutic target for tumors and its underlying mechanisms. Biomed. Pharmacother. 180, 117551 (2024).

29. Hu, S. et al. Prosaposin down-modulation decreases metastatic prostate cancer cell adhesion, migration, and invasion. Mol. Cancer 9, 30 (2010).

30. Ma, Q. et al. Oligodendrocytes drive neuroinflammation and neurodegeneration in Parkinson’s disease via the prosaposin-GPR37-IL-6 axis. Cell Rep. 44, 115266 (2025).

31. Wang, L. et al. A single-cell atlas of glioblastoma evolution under therapy reveals cell-intrinsic and cell-extrinsic therapeutic targets. *Nat*. Cancer 3, 1534–1552 (2022).

32. Yan, T. et al. TGF-β induces GBM mesenchymal transition through upregulation of CLDN4 and nuclear translocation to activate TNF-α/NF-κB signal pathway. Cell Death Dis. 13, 339 (2022).

33. Zhang, G. et al. RBPJ contributes to the malignancy of glioblastoma and induction of proneural-mesenchymal transition via IL-6-STAT3 pathway. Cancer Sci. 111, 4166–4176 (2020).

34. Xie, H. et al. Super-enhancer-driven LIF promotes the mesenchymal transition in glioblastoma by activating ITGB2 signaling feedback in microglia. Neuro. Oncol. 26, 1438–1452 (2024).

35. Milano, M. T. et al. Patterns and timing of recurrence after temozolomide-based chemoradiation for glioblastoma. Int. J. Radiat. Oncol. Biol. Phys. 78, 1147–1155 (2010).

36. Garcia-Diaz, C. et al. Glioblastoma cell fate is differentially regulated by the microenvironments of the tumor bulk and infiltrative margin. Cell Rep. 42, 112472 (2023).

37. de Gooijer, M. C., Guillén Navarro, M., Bernards, R., Wurdinger, T. & van Tellingen, O. An experimenter’s guide to glioblastoma invasion pathways. Trends Mol. Med. 24, 763–780 (2018).

38. Brooks, L. J. et al. The white matter is a pro-differentiative niche for glioblastoma. Nat. Commun. 12, 2184 (2021).

39. Sano, A. et al. Protection by prosaposin against ischemia-induced learning disability and neuronal loss. Biochem. Biophys. Res. Commun. 204, 994–1000 (1994).

40. Jiang, Y. et al. Prosaposin is a biomarker of mesenchymal glioblastoma and regulates mesenchymal transition through the TGF-β1/Smad signaling pathway. J. Pathol. 249, 26–38 (2019).

41. Spitzer, A. et al. Deciphering the longitudinal trajectories of glioblastoma ecosystems by integrative single-cell genomics. Nat. Genet. 57, 1168–1178 (2025).

42. Hart, W. S., Myers, P. J., Purow, B. W. & Lazzara, M. J. Divergent transcriptomic signatures from putative mesenchymal stimuli in glioblastoma cells. Cancer Gene Ther. 31, 851–860 (2024).

43. Halliday, J. et al. In vivo radiation response of proneural glioma characterized by protective p53 transcriptional program and proneural-mesenchymal shift. Proc. Natl. Acad. Sci. U. S. A. 111, 5248–5253 (2014).

44. Jeon, H.-M. et al. Tissue factor is a critical regulator of radiation therapy-induced glioblastoma remodeling. Cancer Cell 41, 1480–1497.e9 (2023).

45. Filbin, M. G. et al. Developmental and oncogenic programs in H3K27M gliomas dissected by single-cell RNA-seq. Science 360, 331–335 (2018).

46. WHO Classification of Tumours Editorial Board. Central Nervous System Tumours. (International Agency for Research on Cancer, 2022).

47. Hao, Y. et al. Integrated analysis of multimodal single-cell data. Cell 184, 3573–3587.e29 (2021).

48. Blanco-Carmona, E. Generating publication ready visualizations for Single Cell transcriptomics using SCpubr. bioRxiv (2022) doi:10.1101/2022.02.28.482303.

49. Timothy Tickle, Itay Tirosh, Christophe Georgescu, Maxwell Brown, Brian Haas. Infercnv. (Bioconductor, 2019). doi:10.18129/B9.BIOC.INFERCNV.

50. Wang, L. et al. The Phenotypes of Proliferating Glioblastoma Cells Reside on a Single Axis of Variation. Cancer Discov. 9, 1708–1719 (2019).

51. McGinnis, C. S., Murrow, L. M. & Gartner, Z. J. DoubletFinder: Doublet detection in single-cell RNA sequencing data using artificial nearest neighbors. Cell Syst. 8, 329–337.e4 (2019).

52. Andreatta, M. & Carmona, S. J. UCell: Robust and scalable single-cell gene signature scoring. Comput. Struct. Biotechnol. J. 19, 3796–3798 (2021).

53. Ianevski, A., Giri, A. K. & Aittokallio, T. Fully-automated and ultra-fast cell-type identification using specific marker combinations from single-cell transcriptomic data. Nat. Commun. 13, 1246 (2022).

54. Yu, G., Wang, L.-G., Han, Y. & He, Q.-Y. clusterProfiler: an R package for comparing biological themes among gene clusters. OMICS 16, 284–287 (2012).

