## Supplementary Information for "Unveiling Oligodendrocyte-Lineage Differentiation in the Glioblastoma Infiltrative Zone Through Spatial Transcriptomics"

**Table S1. Patient demographics and clinical information for the stRNA-seq and IHC cohort.**

| Sample Number | Gender | Gliolan | Diagnosis | IDH status | MGMT | MKI67 | ATRX | TP53 |
|---|---|---|---|---|---|---|---|---|
| PT#1 | M | Yes | Glioblastoma grade IV | WT | NT | 25% | WT | +/- |
| PT#2 | M | Yes | Glioblastoma grade IV | WT | Unmet | 25% | NT | 80% |
| PT#3 | M | No | Giant-cell Glioblastoma grade IV | WT | Unmet | 20% | NT | 90% |

WT= Wild-type, NT= Non-tested, Unmet= Unmethylated. MKI67 and TP53 are expressed as the percentage of MKI67-positive cells and TP53-mutated cells.

**Table S2. Sample IDs of analyzed stRNA-seq samples.**

| Patient | Area-Sample ID |
|---|---|
| PT#1 | Inner edge of the tumor-22SC1A |
| PT#2 | Core of the tumor-22SC1B |
| PT#3 | Outer edge of the tumor-22SC1C |

**Table S3. Patient demographics and clinical information for the FISH cohort.**

| Sample Number | Gender | Gliolan | Diagnosis | IDH status | MGMT | MKI67 | ATRX | TP53 |
|---|---|---|---|---|---|---|---|---|
| PT#1 | M | Yes | Glioblastoma grade IV | WT | Unmet | 25% | NT | 80% |
| PT#4 | M | Yes | Glioblastoma grade IV | WT | Unmet | 20% | NT | 90% |
| PT#5 | F | No | Glioblastoma grade IV | WT | Met | 20% | NT | 40% |
| PT#6 | M | Yes | Glioblastoma grade IV | WT | NT | 25% | WT | 40% |
| PT#7 | F | Yes | Glioblastoma grade IV | WT | Met | 25% | WT | WT |

WT= Wild-type, NT= Non-tested, Unmet= Unmethylated. MKI67 and TP53 are expressed as the percentage of MKI67-positive cells and TP53-mutated cells.

**Table S3. Patient demographics and clinical information for the FISH cohort**

Provided as .xlsx files.

**Table S5. CellChat ligand-receptor analysis table of spots in the outer edge of the tumor.**

Provided as .xlsx files.

**Fig. S1: Data quality assessment and CNV inference.**

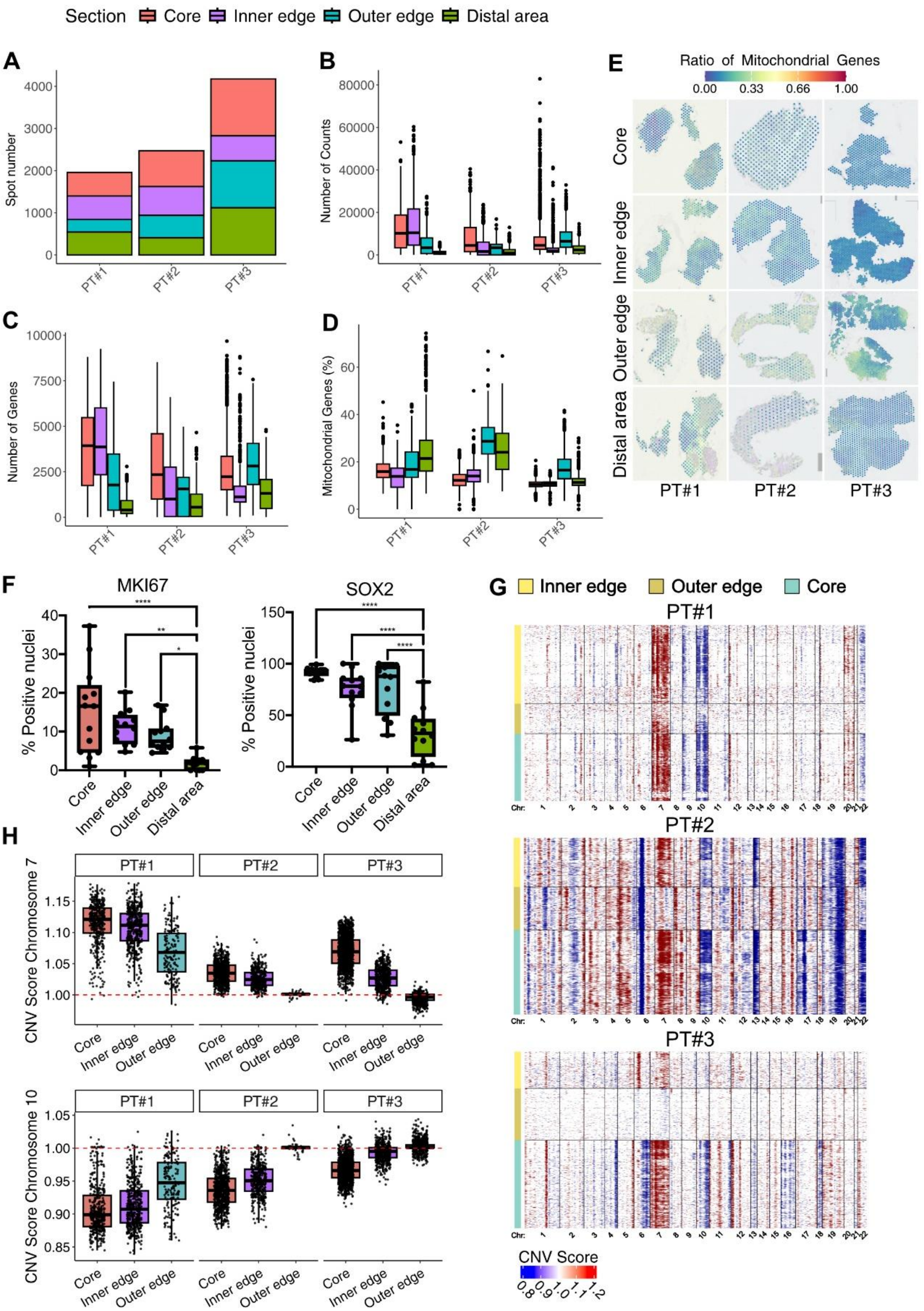
Section Core Inner edge Outer edge Distal area
A
Spot number
PT#1
PT#2
PT#3
B
Number of Counts
C
Number of Genes
D
Mitochondrial Genes (%)
E
Ratio of Mitochondrial Genes
0.00 0.33 0.66 1.00
Core
Inner edge
Outer edge
Distal area
F
MKI67
SOX2
% Positive nuclei
Core
Inner edge
Outer edge
Distal area
G
Inner edge Outer edge Core
H
CNV Score Chromosome 7
CNV Score Chromosome 10
CNV Score
0.8 0.9 1.0 1.1 1.2

**(A)** Stacked barplot showing the number of spots per area in the three patients included in the study. **(B,C,D)** Boxplots representing quality control metrics for the four sampled areas of the three patients. Metrics used for spot quality assessment included the number of transcripts per spot (**B**), the number of expressed genes per spot (**C**), and the percentage of expressed mitochondrial genes per spot (**D**). (**E**) Spatial feature plot showing the percentage of expressed mitochondrial genes per spot in the four areas of the three patients. **(F)** Boxplot of MKI67 (left) and SOX2 (right) positive cells in near-adjacent tissue sections from the four areas of the three patients. For each area, we quantified three frames and normalized the number of positive cells on the total number of DAPI positive nuclei. One way ANOVA test with *post-hoc* Tuckey's test. *p < 0.05, **p < 0.01, ***p < 0.001, ****p < 0.0001. **(G)** Heatmap of CNVs across each genomic position (columns) for each spot (rows) in the three different patients. CNVs were inferred from the transcriptome of single spots using the InferCNV tool, using spots in the distal area as reference for the analysis. In the heatmaps, spots are grouped by sampled area. **(H)** Boxplots of CNV scores for chromosome 7 (upper) and chromosome 10 (lower) for spots in the different areas, the distal area was used as a reference. The red dotted line indicates the baseline CNV score of 1, representing diploidy.

**Fig. S2: Generation of an integrated scRNA-seq dataset and nontumor spot analysis.**

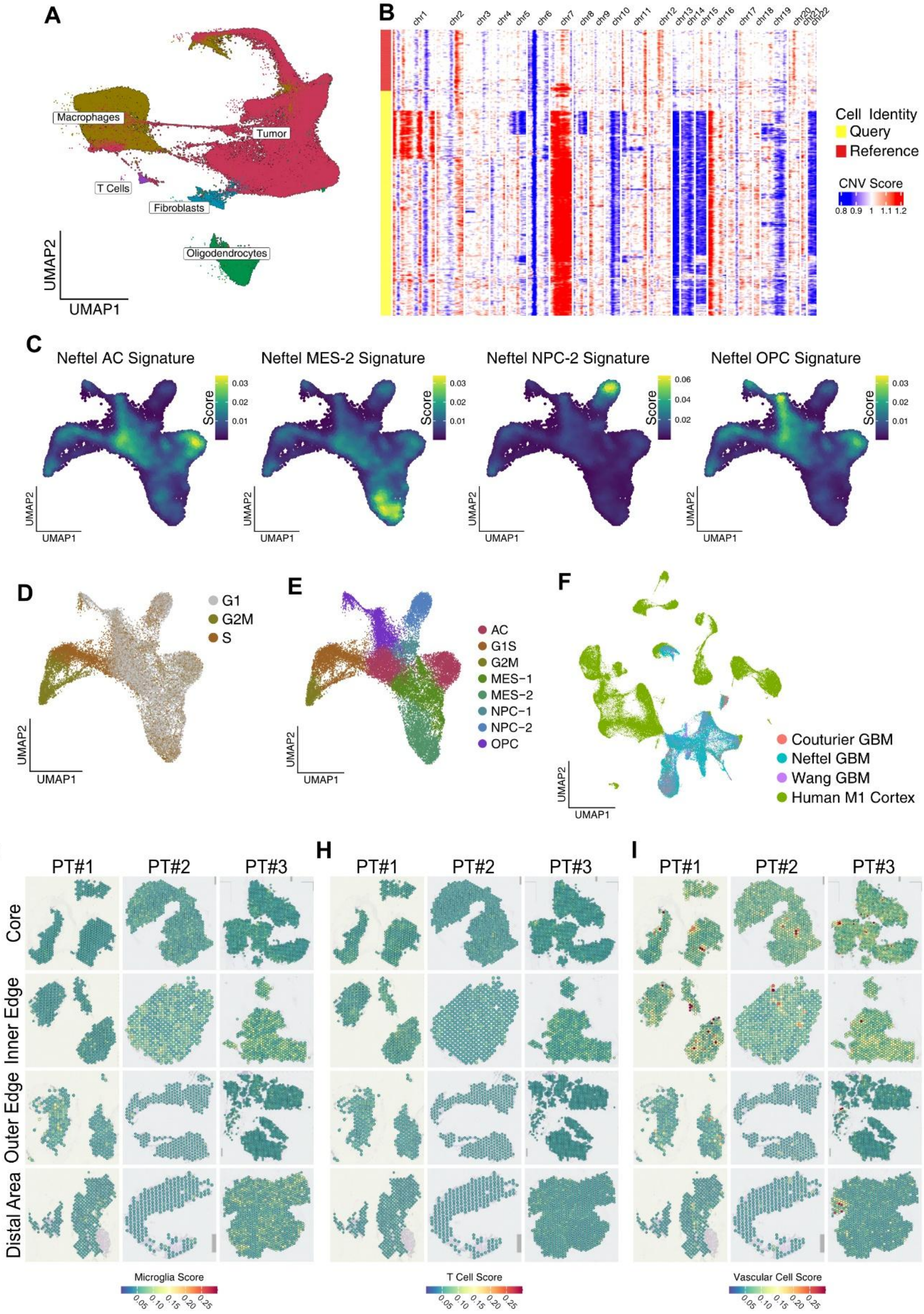
A
Macrophages
Tumor
T Cells
Fibroblasts
Oligodendrocytes
UMAP2
UMAP1
B
chr1 chr2 chr3 chr4 chr5 chr6 chr7 chr8 chr9 chr10 chr11 chr12 chr13 chr14 chr15 chr16 chr17 chr18 chr19 chr20 chr21 chr22
Cell Identity
Query
Reference
CNV Score
0.8 0.9 1 1.1 1.2
C
Neftel AC Signature
Neftel MES-2 Signature
Neftel NPC-2 Signature
Neftel OPC Signature
Score
D
G1
G2M
S
E
AC
G1S
G2M
MES−1
MES−2
NPC−1
NPC−2
OPC
F
Couturier GBM
Neftel GBM
Wang GBM
Human M1 Cortex
G
H
I
PT#1
PT#2
PT#3
Core
Inner Edge
Outer Edge
Distal Area
Microglia Score
T Cell Score
Vascular Cell Score
0.05 0.10 0.15 0.20 0.25

**(A)** UMAP projection of all integrated scRNA-seq GBM datasets, colors indicate the annotated cell type. Nontumor cell subtypes were annotated based on clustering and expression of published lists of marker genes. **(B)** Heatmap of CNVs across each genomic position for each cell in the GBM scRNA-seq datasets. CNVs were inferred from the transcriptome of single cells using the InferCNV tool. The red cluster in the top part of the heatmap represents cells used as reference for the CNV analysis. **(C)** Density plots showing the expression scores of Neftel's GBM state gene modules in tumor cells of the GBM reference dataset. **(D)** UMAP plots of tumor cells of the GBM reference scRNA-seq datasets grouped by cell cycle phase. **(E)** Annotated UMAP plot of tumor cells of the GBM reference scRNA-seq datasets. **(F)** Annotated UMAP plot of the reference scRNA-seq dataset used to study spatial spot composition. The reference was obtained by integrating several published GBM samples to an annotated public human motor cortex scRNA-seq dataset. Cells are colored by the dataset of origin. **(G,H,I)** Surface plots of microglia (**G**), T cell (**H**), and vascular cell (**I**) marker expression scores in all spatial spots for all samples.

**Fig. S3: Analysis of tumor spot transcription patterns and signaling networks.**

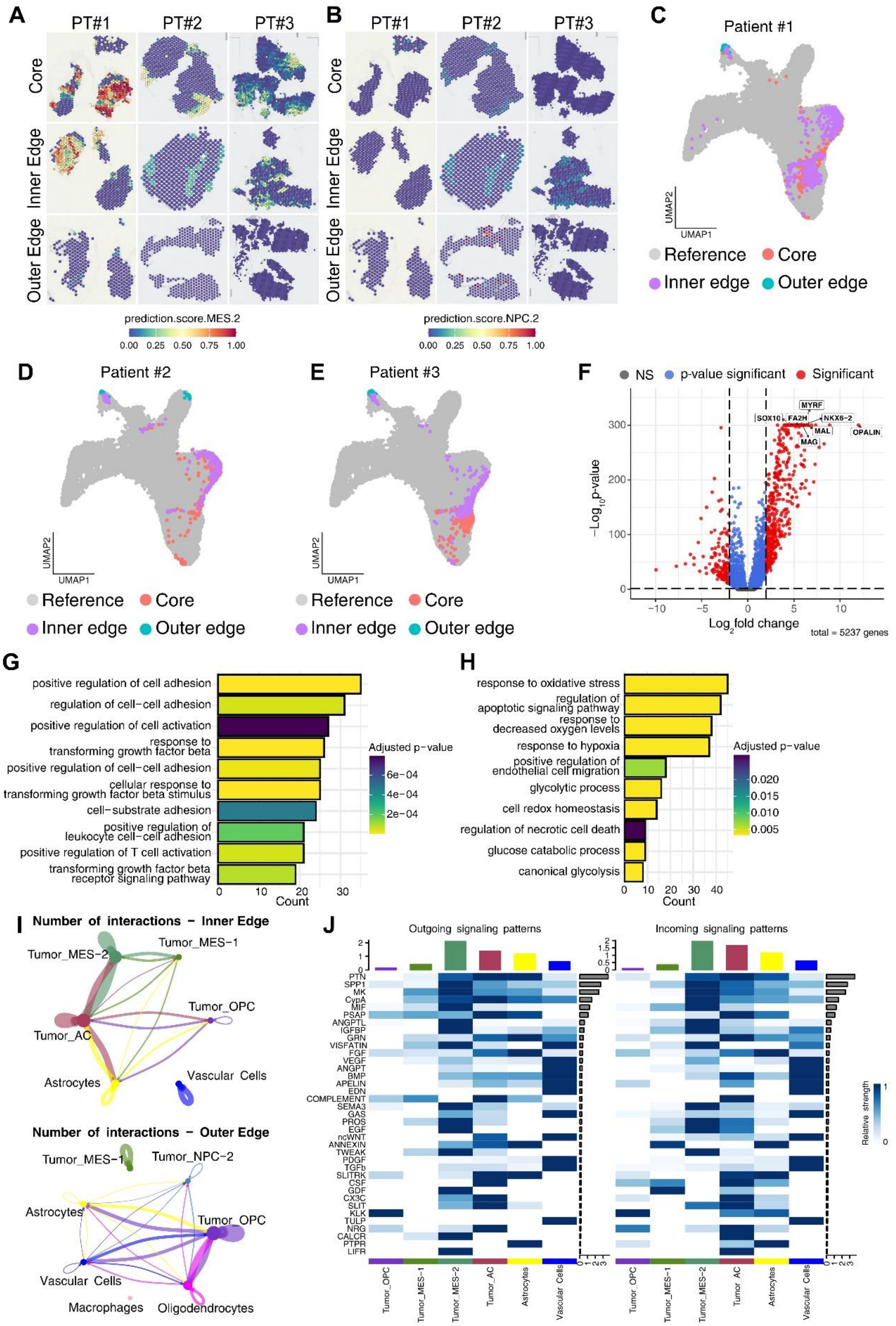
A
PT#1
PT#2
PT#3
Core
Inner Edge
Outer Edge
prediction.score.MES.2
0.00 0.25 0.50 0.75 1.00
B
PT#1
PT#2
PT#3
Core
Inner Edge
Outer Edge
prediction.score.NPC.2
0.00 0.25 0.50 0.75 1.00
C
Patient #1
UMAP2
UMAP1
Reference
Core
Inner edge
Outer edge
D
Patient #2
E
Patient #3
F
NS
p-value significant
Significant
MYRF
SOX10
FA2H
NKX6-2
MAL
MAG
OPALIN
−Log10 p-value
Log2 fold change
total = 5237 genes
G
positive regulation of cell adhesion
regulation of cell-cell adhesion
positive regulation of cell activation
response to transforming growth factor beta
positive regulation of cell-cell adhesion
cellular response to transforming growth factor beta stimulus
cell-substrate adhesion
positive regulation of leukocyte cell-cell adhesion
positive regulation of T cell activation
transforming growth factor beta receptor signaling pathway
Count
Adjusted p-value
H
response to oxidative stress
regulation of apoptotic signaling pathway
response to decreased oxygen levels
response to hypoxia
positive regulation of endothelial cell migration
glycolytic process
cell redox homeostasis
regulation of necrotic cell death
glucose catabolic process
canonical glycolysis
Count
Adjusted p-value
I
Number of interactions − Inner Edge
Tumor_MES-2
Tumor_MES-1
Tumor_OPC
Tumor_AC
Vascular Cells
Astrocytes
Number of interactions − Outer Edge
Tumor_MES-1
Tumor_NPC-2
Astrocytes
Tumor_OPC
Vascular Cells
Macrophages
Oligodendrocytes
J
Outgoing signaling patterns
Incoming signaling patterns
Relative strength
Tumor_OPC
Tumor_MES-1
Tumor_MES-2
Tumor_AC
Astrocytes
Vascular Cells

**(A,B)** Surface plots of MES-2 (**A**), and NPC-2 (**B**), cell state prediction score in tumor spatial spots for all samples in the three tumor areas. The score represents the confidence of the predicted cell state on a scale from 0 to 1, based on the number of anchor genes found between the query spatial spot and the scRNA-seq reference embedding. **(C,D,E)** UMAP plots showing the mapping of tumor spot projection for the three tumor areas of patient 1 (**C**), 2 (**D**), and 3 (**E**) on the reference GBM scRNA-seq embedding. **(F)** Volcano plot of the differential expression analysis of tumor spots in the outer edge of the tumor versus tumor spots in the core and inner edge of the tumor. **(G)** Barplot showing representative terms of the gene ontology enrichment analysis performed on genes differentially expressed by tumor spots in the inner edge of the tumor versus tumor spots in the other areas. **(H)** Barplot showing representative terms of the gene ontology enrichment analysis performed on genes differentially expressed by tumor spots in the core of the tumor versus tumor spots in the other areas. **(I)** Circle plots showing the aggregated signaling networks for the inner edge (top) and the outer edge (bottom) of the tumor. The size of the dot is proportional to the numerosity of that spot type in the data, while the size of the arrow is proportional to the number of predicted interactions between the two involved cell types. Arrows and dots are colored by spot type. **(J)** CellChat heatmaps showing the relative signaling strength of the most active pathways in the different spot types of the inner edge of the tumor. The top histogram shows the total signaling strength of a cell type by summing the contribution of all of the pathways shown. The right histogram shows the total strength of a pathway by summing the strengths of all cell types shown. The left heatmap shows outward interactions, while the right heatmap shows inward interactions.

**Fig. S4: Validation of the prevalent OPC-like nature of infiltrating GBM cells.**

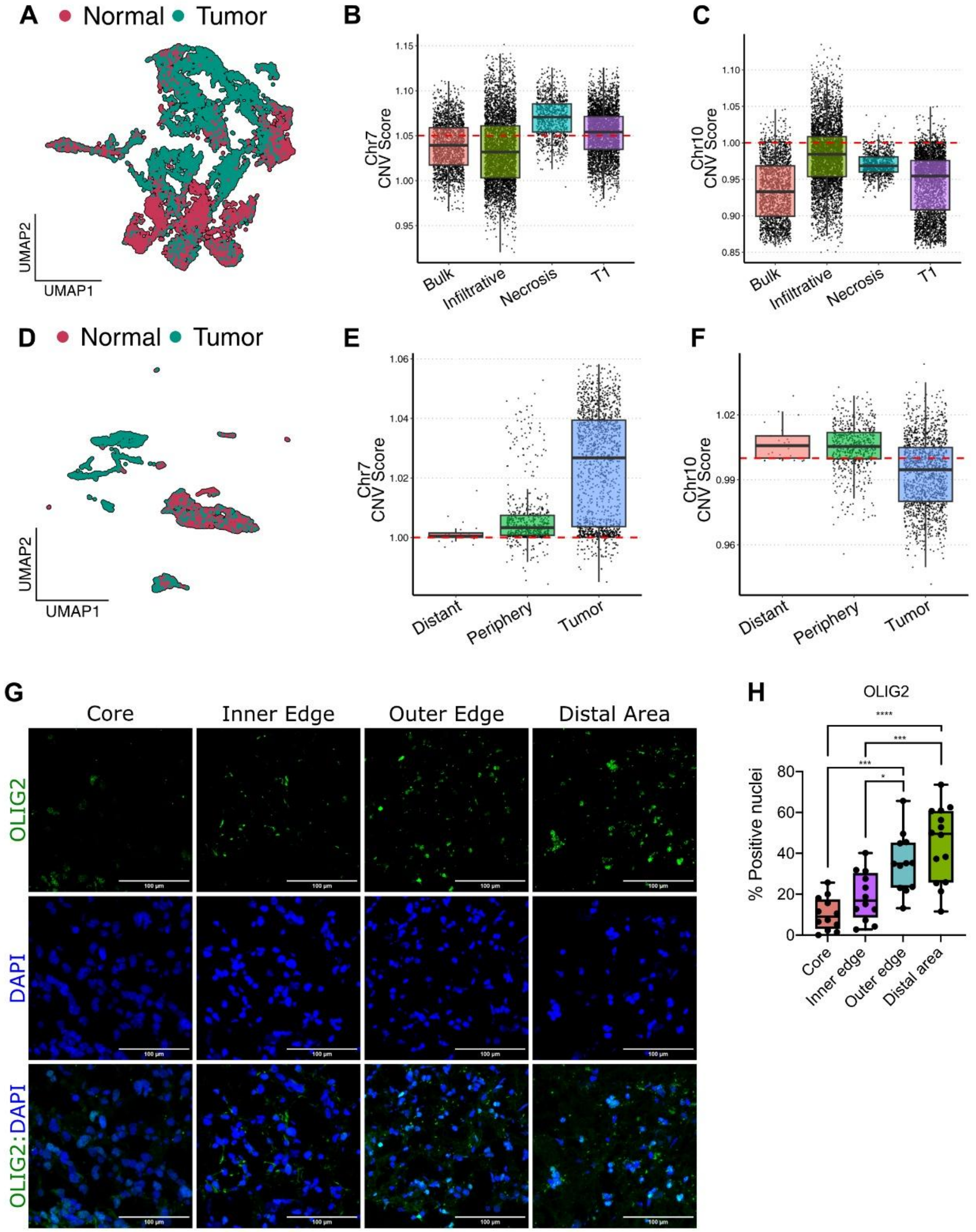


**(A)** UMAP plot of all integrated spots from the Greenwald et al. spatial transcriptomics dataset. Spots are colored by tumor/normal annotation based on the CNV analysis. **(B,C)** Boxplots of CNV scores for chromosome 7 (**B**) and chromosome 10 (**C**) for spots in the different areas of the

Greenwald et al. spatial transcriptomics dataset. The red dotted line indicates the CNV score thresholds used for tumor/normal annotation. **(D)** UMAP plot of all integrated cells from the Darmanis et al. scRNA-seq dataset. Cells are colored by tumor/normal annotation based on the CNV analysis. **(E,F)** Boxplots of CNV scores for chromosome 7 (**E**) and chromosome 10 (**F**) for cells in the different areas of the Darmanis et al. scRNA-seq dataset. The red dotted line indicates the CNV score thresholds used for tumor/normal annotation. **(G)** Representative OLIG2 IHC images of tissue sections from the four sampled areas. **(H)** Boxplot of OLIG2 positive cells in near-adjacent tissue sections from the four areas of the three patients. For each area, we quantified three frames and normalised the number of positive cells on the total number of DAPI positive nuclei. One way ANOVA test with *post-hoc* Tuckey's test. *p < 0.05, **p < 0.01, ***p < 0.001, ****p < 0.0001.
